# Identification of activators of human fumarate hydratase by quantitative high-throughput screening

**DOI:** 10.1101/690909

**Authors:** Hu Zhu, Olivia W. Lee, Pranav Shah, Ajit Jadhav, Xin Xu, Samarjit Patnaik, Min Shen, Matthew D. Hall

## Abstract

Fumarate hydratase (FH) is a metabolic enzyme that is part of the Krebs-cycle, and reversibly catalyzes the hydration of fumarate to malate. Mutations of the *FH* gene have been associated with fumarate hydratase deficiency (FHD), hereditary leiomyomatosis, renal cell cancer (HLRCC), and other diseases. Currently there are no high-quality small molecule probes for studying human fumarate hydratase. To address this, we developed a quantitative high throughput screening (qHTS) FH assay and screened a total of 57,037 compounds from in-house libraries in dose-response. While no inhibitors of FH were confirmed, a series of phenyl-pyrrolo-pyrimidine-diones were identified as activators of human fumarate hydratase. These compounds were not substrates of fumarate hydratase, were inactive in a malate dehydrogenase counter screen, and showed no detectable reduction–oxidation activity. The binding of two compounds from the series to human fumarate hydratase was confirmed by microscale thermophoresis. The low hit rate in this screening campaign confirmed that FH is a ‘tough target’ to modulate, and the small molecule activators of human fumarate hydratase reported here may serve as a starting point for further optimization and development into cellular probes of human FH and potential drug candidates.

## Introduction

Fumarate hydratase (FH) is an important metabolic enzyme in the Krebs-cycle, which reversibly catalyzes the conversion of fumarate to malate (hydration). It is encoded by the *FH* gene, which produces two isoforms of protein by alternative splicing, a cytosolic form and a mitochondrial form. The mitochondrial form of FH contains a N-terminal mitochondrial targeting sequence, which is removed in the mitochondrion to generate the same protein as that in the cytoplasm. Despite decades of research, a detailed understanding of the mechanism of FH’s biochemical activity remains a matter of debate. The active enzyme is a homo-tetramer in which each active site is made up of residues at the interface of three of the four subunits^1^. FH participates in ATP production in mitochondria and the metabolism of amino acids and fumarate in cytoplasm.

In humans, mutations of *FH* gene lead to defects of the enzymatic activity either by indirectly compromising the integrity of the protein’s core architecture or directly affecting residues within the active site that regulate catalytic activity of the enzyme^2^. Mutations of *FH* gene have been mainly associated with two heritable diseases: fumarate hydratase deficiency, and hereditary leiomyomatosis and renal cell cancer (HLRCC). Homozygous germline mutations in *FH* gene cause autosomal recessive fumarate hydratase deficiency syndrome. It is characterized by early-onset hypotonia, profound psychomotor retardation, seizures, facial dysmorphism and brain abnormalities^3^. Mutations of *FH* gene severely impair the activity of enzyme which leads to the defect of energy production and the accumulation of fumarate, which is believed to be the cause of clinical symptoms^4^. Heterozygous germline mutations in *FH* gene predispose individuals to HLRCC, characterized with benign leiomyomas of the skin and the uterus and early-onset of type II papillary renal cell carcinoma^5^. In affected individuals, loss of heterozygosity (LOH) in the wild-type allele by somatic mutations leads to severe reduction or absence of fumarate hydratase activity, which is followed by the increased intracellular level of fumarate. Accumulated fumarate acts as an oncometabolite to induce tumorigenesis through competitively inhibit α-ketoglutarate–dependent dioxygenase enzymes and post-translationally modify proteins by succination^6, 7^. Aside from FH deficiency and HLRCC, studies have also shown that fumarate hydratase is involved in hypertension, type 2 diabetes and diabetic kidney disease^8–10^. For example, deletion of FH in mouse pancreatic β cells leads to progressive glucose intolerance and diabetes after 6-8 weeks. In human, the fumarate level is higher in islets from donors of type 2 diabetes than the normal donors, and high glucose produce no further increase of fumarate^10^.

Currently there are no biochemically characterized inhibitors of FH, and no known activator for this enzyme. Two series of inhibitors have been reported to inhibit the activity of human^11^ or *Mycobacterium tuberculosis* FH^12^. In an attempt to identify nutrient-dependent cytotoxic compounds, Takeuchi *et al*^11^ screened a collection of 6,000 small molecules and discovered a new class of inhibitors of human FH (compound **1-3** with pyrrolidinone structure). In another report, Kasbekar^12^ reported that two inhibitors (compound **7** and **8**) were identified for the *Mycobacterium tuberculosis* fumarate hydratase (tbFH) after screening a collection of 479,984 small molecules.

To identify small molecule activators or inhibitors of human FH, we developed a fluorescence-based FH assay by coupling to malate dehydrogenase (MDH)/diaphorase/resazurin and optimized the assay for quantitative high-throughput screening (qHTS) in 1536-well plates. We proceeded to screen a collection of small molecule libraries to identify new activators or inhibitors of human FH. A total of 57,037 small molecular compounds were screened, and majority of hits were triaged by malate dehydrogenase (MDH) counter screen assay. One series of the activators for human fumarate hydratase was confirmed and validated in the current study.

## Materials and methods

### Reagents and chemical libraries

Recombinant human fumarate hydratase (with 6 histidine tag in the N-terminal of protein), dithiothreitol (DTT) and HBSS solution were purchased from Thermofisher Scientific (Waltham, MA). L-malate dehydrogenase from pig heart, diaphorase from *clostridium kluyveri*, (β-NAD (NAD+), resazurin, Brij 35, sodium fumarate dibasic, malic acid and horse radish peroxidase (HRP) were purchased from Sigma (St. Louis, MO). Fumarate hydratase-IN-2 was purchased from MedChem Express (Monmouth Junction, NJ). Compound 8 was purchase from Enamine (Monmouth Junction, NJ). Amplex Red (10-Acetyl-3,7-dihydroxyphenoxazine) was purchased from Cayman Chemical (Ann Arbor, MI). Six libraries were screened, including Sytravon (a retired Pharma screening collection that contains a diversity of novel small molecules, NCATS), NPACT (NCATS Pharmacologically Active Chemical Toolbox, NCATS), NPC (NCATS Pharmaceutical Collection, NCATS),^13^ MIPE (Mechanism Interrogation Plate, NCATS), Kinase inhibitor library (an in-house collection of kinase inhibitors, NCATS) and LOPAC (1,280 Pharmacologically Active Compounds Sigma-Aldrich, St. Louis, MO) libraries.

### High-throughput screen and counter screen

For primary high-throughput screen, 3 μl of FH solution (containing 13.33nM human fumarate hydratase, 13.33IU/ml malic dehydrogenase, 0.2mM NAD, 0.067mg/ml diaphorase and 0. 067mM resazurin in the assay buffer (50 mM Tris pH 8.0, 5 mM MgCl2, 0.01% Brij 3) was dispensed into each well in a black solid bottom 1536-well assay plate (Greiner Bio-One) using a BioRAPTR FRD dispenser (Beckman Coulter, Brea, CA). A 1536-well pintool dispenser outfitted with 20 nl pins (Wako Automation, San Diego, CA) was used to transfer 20 nL of DMSO-solubilized compound (Cherrypick plates) to each 1536-well assay plate. Each compound was screened at five concentrations: 15 nM, 151 nM, 1.5 μM, 15.4 μM, 76.9 μM. Following compound transfer, plates were incubated in room temperature for 10 minutes. 1 μL of substrate solution containing fumaric acid (160 μM) was dispensed via BioRAPTR FRD to initiate the reaction. Plates were immediately transferred to a ViewLux microplate imager (PerkinElmer, Waltham, MA), and any resulting resorufin fluorescence was measured (exitation/emission, 525/598 nm) at 0 and 15 minutes. The exposure time is 1 second. Fluorescence from each well was normalized using enzyme-free and DMSO-treated control wells on each plate, and changes in fluorescence (ΔRFU) were calculated using the difference in fluorescent signal for each well at 15 minutes versus 0 minutes.

The malate dehydrogenase counter screen was adapted from the primary high-throughput screen protocol. 3 μl MDH solution (containing 13.33IU/ml malic dehydrogenase, 0.2mM NAD, 0. 067mg/ml diaphorase and 0.067mM resazurin in assay buffer (50 mM Tris pH 8.0, 5 mM MgCl2, 0.01% Brij 3)) was dispensed in each well in the 1536-well plate. Plates were incubated at room temperature for 10 minutes after compound transfer. 1 μL of substrate solution containing malic acid (160uM) was dispensed to initiate the reaction. Fluorescence was measured (ex540, em590 nm) at 0 and 5 minutes. The fluorescence was normalized using enzyme-free and DMSO-treated control wells on each plate, and ΔRFU was calculated using the difference in fluorescent signal for each well at 5 minutes versus 0 minutes.

### REDOX assay

Amplex red assay was used to detect the REDOX activity of small molecules. Briefly, 2.5 μl HBSS solution (1.26 mM CaCl2, 0.49 mM MgCl2, 1 g/L D-glucose) was dispensed into each well in a black solid bottom 1536-well assay plate. 20 nl of compounds was pinned into the plate and the background fluorescence (RFU_0min_) was measured in the Viewlux (excitation/emission, 525/598 nm). 2.5 μl freshly made 2X Amplex Red solution (100 μM Amplex Red, 200 μM DTT and 2 U/mL HRP in 1xHBSS solution) was added to each well. After 15min incubation in room temperature, fluorescence (RFU_15min_) was measured in the Viewlux with same setting. REDOX activity of each compound was calculated using corrected fluorescence values (ΔRFU=RFU_15min_-RFU_0min_). Negative control is DMSO, and positive controls are walrycin B and chlorinal.

### Microscale thermophoresis

Recombinant human fumarate hydratase, which contains a polyhistidine tag at the C-terminus, was labeled with Monolith His-Tag Labeling Kit RED-tris-NTA kit (NanoTemper Technologies, München, Germany). Briefly, the protein was diluted to 200 nM in PBS-T buffer (137 mM NaCl, 2.5 mM KCl, 10 mM Na2HPO4, 2 mM KH2PO4, pH 7.4;0.05 % Tween-20). The RED-tris-NTA dye was also diluted in PBS-T to 100 nM. The diluted protein and dye were mixed in 1:1 volume ratio and incubated for 30 min at room temperature (RT). After centrifuge 15,000 x g for 10 minutes at 4°C, the supernatant was transferred to a new tube and ready to use.

Compounds were serially diluted in DMSO (16 points, 1:2 dilutions, from 10mM), and then further diluted 20 times in PBS-T buffer. 10ul labeled protein and 10ul diluted compounds were mixed and incubate at RT for 2-3 minutes. The samples were loaded in the standard treated capillaries and measured in a NanoTemper Monolith NT.115 LabelFree instrument (NanoTemper Technologies, München, Germany). The samples were measured at LED power of 40%, and MST power of 40 % and 60 %. A laser-on time of 30 s and a laser-off time of 5 s was used in this experiment. Curve fitting was performed using GraphPad Prism 5 (La Jolla, CA).

### *In vitro* ADME Studies

The physicochemical and pharmacokinetics properties of compounds were measured using the following high-throughput *in vitro* assays: rat microsomal stability assay, parallel artificial membrane permeability assay (PAMPA) and aqueous kinetic solubility assays. The details of these assays are described in a previous study^14^.

### qHTS Data Analysis

Analysis of compound concentration–response data was performed as previously described^15^. Briefly, raw plate reads for each titration point were first normalized relative to the no enzyme control and DMSO-only wells as follows: % Activity = ((V_compound_ – V_DMSO_)/(V_DMSO_ – V_no enzyme_)) × 100, where V_compound_ denotes the compound well values, Vno enzyme denotes the median value of the no enzyme control wells, and V_DMSO_ denotes the median values of the DMSO-only wells, and then corrected by applying a NCGC in-house pattern correction algorithm using compound-free control plates (i.e., DMSO-only plates) at the beginning and end of the compound plate stack^16^. Concentration–response titration points for each compound were fitted to a four-parameter Hill equation^17^ yielding concentrations of half-maximal activity (AC_50_) and maximal response (efficacy) values. Compounds were designated as Class 1–4 according to the type of concentration–response curve observed^15, 18^. Curve classes are heuristic measures of data confidence, classifying concentration–responses on the basis of efficacy, the number of data points observed above background activity, and the quality of fit. Compounds with curve classes 1.1, 1.2, 2.1, 2.2 (activators) and −1.1, −1.2, −2.1 or −2.2 (inhibitors) were considered active. Class 4 compounds were considered inactive. Compounds with other curve classes were deemed inconclusive. Active compounds were cherrypicked for follow up experiments.

## Results

### Assay design, optimization and miniaturization

To identify activators or inhibitors of human FH, we developed a fluorescence-based FH assay, adapted from a previous study^12^. In our assay format, the FH reaction is coupled to two sequential enzymatic reactions: MDH and diaphorase reactions. In the first coupled MDH reaction, L-malate produced by FH is converted into oxaloacetate, removing L-malate from the system and driving the FH forward reaction. The reduced form of nicotinamide adenine dinucleotide (NADH) generated by MDH is utilized in the second coupled reaction by diaphorase to convert resazurin to resorufin (Fig 1A), with the further benefit of regenerating NAD+ to drive the MDH biochemical reaction. Resorufin is strongly fluorescent, with an excitation wavelength at 530-540 nm and emission at 585-595 nm^19, 20^. Therefore, FH activity can be evaluated by measuring fluorescence signal of resorufin.

**Figure 1.**
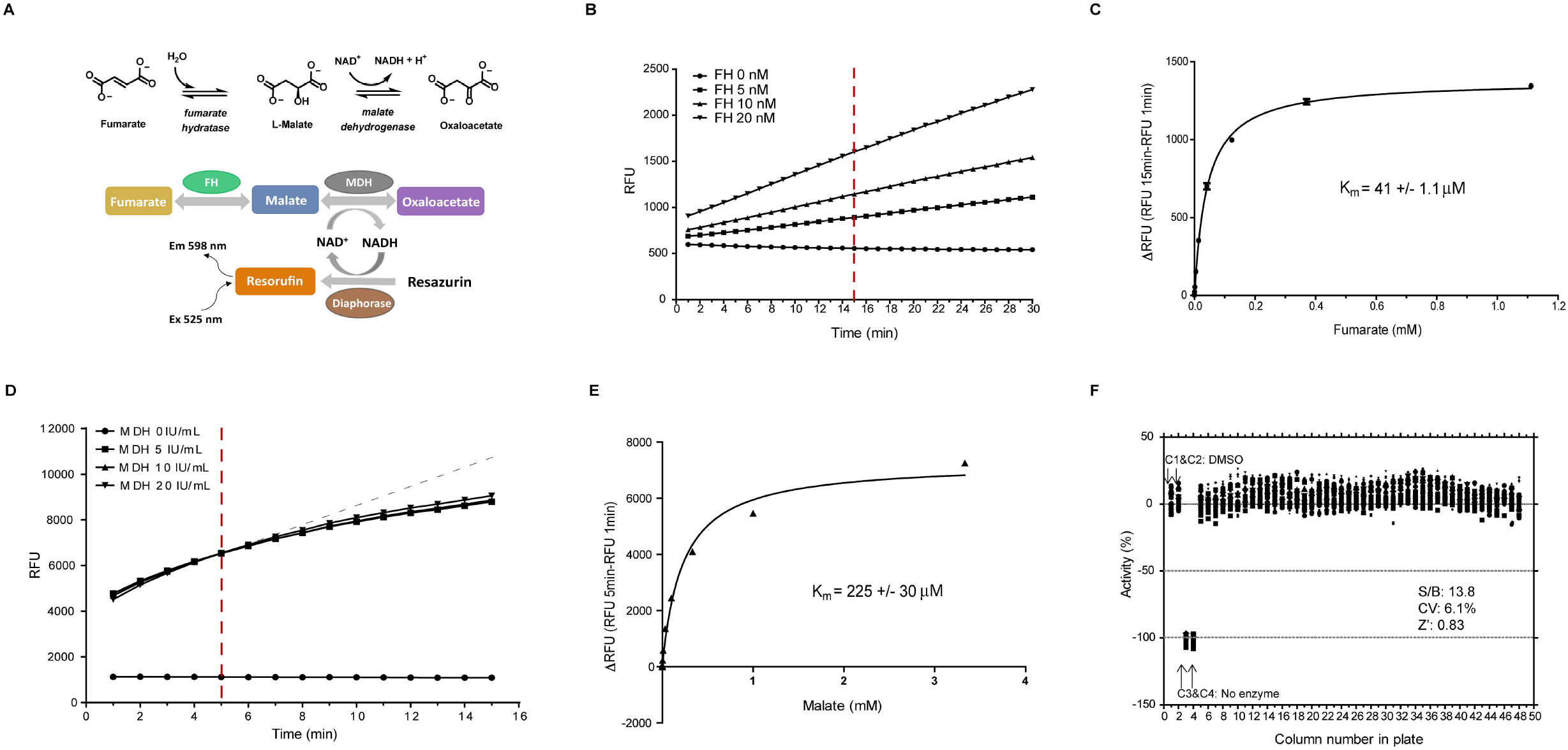
Validation of FH primary assay and MDH counter assay in qHTS screen. **A**. Schematic diagram of the fluorescence-based high-throughput screening assay used to evaluate fumarate hydratase activity. FH enzymatic reaction is coupled to MDH and diaphorase reaction for the detection of resorufin fluorescence. **B**. Time course of FH assay with varying concentration of FH enzyme (0, 5, 10, 20 nM). Fluorescence was continuously measured every minute for 30 minutes. The red dot line indicates the time the signals were measured. **C**. Determine Km value of fumarate for FH enzyme. Reactions were assembled with 10 nM FH, indicated concentration of fumarate, and incubated at room temperature for 15min. D. Time course of MDH assay with varying concentration of MDH enzyme (0, 5, 10, 20 IU/ml). Fluorescence was continuously measured every minute for 30 minutes. The red dot line indicates the time the signals were measured. **E**. Determine Km value of malate for MDH enzyme. Reactions were assembled with 10 IU/ml MDH, indicated concentration of malate, and incubated at room temperature for 5min. **F**. Scatter plot of the percentage of FH activity from a 1536-well DMSO plate to validate the assay. Column 1 and 2 are DMSO neutral control, and column 3 and 4 were no FH enzyme control. In this plate, the ratio of signal/background (S/B) was 13.8, coefficient of variation (CV) was 6.1, and Z factor was 8.3.

To optimize assay conditions, the initial linear velocity of the enzymatic reaction was determined in the presence of excess fumarate. We found that the enzymatic reaction was linear over 30 minutes at concentration of up to 20 nM FH enzyme (Fig 1b). We selected 10 nM FH enzyme concentration and monitored the reaction for 15 min at room temperature (incubations not amenable to a high-throughput platform) for remaining experiments. The K_m_ value of fumarate was determined as 41 ± 1.1 μM by monitoring reaction kinetics over a wide range of fumarate concentrations (Fig 1C). As we sought to identify both activators and inhibitors, the FH assay was performed with a substrate concentration near the K_m_ value of fumarate (40μM). A known liability of coupled enzymatic assays is that small molecules appearing to be active from HTS may in fact be modulating the activity of MDH or diaphorase, rather than the target itself. To enable the triage of false positives, we developed a MDH counter-assay, omitting FH and adding the MDH substrate malate to initiate the reaction. The kinetics of the MDH counter assay were evaluated at variable concentrations of MDH enzyme and at several time points. The biochemical reaction was found to be in the linear range for the first 5-6 min at concentrations up to 20 IU/mL MDH enzyme (Fig 1D). The Km value of malate was determined by MDH assay as 225 ± 30 μM (Fig 1E). This concentration of malate was selected for the MDH counter-assay.

To assess the reproducibility and robustness of the assay, FH assay was performed in white solid bottom 1536-well Greiner microplates and statistics was analyzed. Buffer containing enzyme was dispensed into a microplate from columns 1, 2 and 5-48. The reagent without FH enzyme was dispensed into columns 3 and 4 as an enzyme-free negative control. DMSO (23 nL) was transferred into all wells, and then incubated with enzyme for 10 minutes. The reaction was then initiated by adding fumarate. The kinetics of reaction was measured at t = 0 min and t = 15 min (Table 1). As shown in Figure 1F, signal-to-background (S/B) was 13.8, coefficient of variation was 6.1% and Z’ factor was 0.83. The assay conditions result in less than 10% substrate conversion during the course of the assay. To ensure the assay is amenable to storage of reagents over the multiple hours that a qHTS assay would require to be initiated, enzymatic activity was assessed over storage times. FH was found to be unstable when stored at room temperature, rapidly losing activity within 2 h, but was quite stable when stored on ice for at least 6 hours (Supplemental Fig 1).

**Table 1.** Protocol of FH assay and MDH counter-assay.

Protocol of FH assay
| Step | Parameter | Value | Descriptoin |
| --- | --- | --- | --- |
| 1 | Reagent a | 3 μl | Dispense reagent a: 13.33 nM fumarate hydratase, 13.33 units/mL MDH, 0.2 mM NAD<sup>+</sup>, 0.067 mg/mL diaphorase, 0.067 mM resazurin (in 50 mM Tris pH 8, 5 mM MgCl<sub>2</sub>, 0.01% Brij 35). |
| 2 | Reagent b | 3 μl | Dispense reagent b: 13.33 units/mL MDH, 0.2 mM NAD<sup>+</sup>, 0.067 mg/mL diaphorase, 0.067 mM resazurin (in 50 mM Tris pH 8, 5 mM MgCl<sub>2</sub>, 0.01% Brij 35). |
| 3 | Compounds | 23 nL | Pin transfer of compounds |
| 4 | Time | 10 min | Incubate at room temperature |
| 5 | Reagent | 1 μl | Dispense 1 μl 160 μM fumaric acid |
| 6 | Read | 525 nm / 598 nm | ViewLux read twice at 15 minutes interval |

Protocol of MDH counter-assay
| Step | Parameter | Value | Descriptoin |
| --- | --- | --- | --- |
| 1 | Reagent a | 3 μl | Dispense reagent a: 13.33 units/mL MDH, 0.2 mM NAD<sup>+</sup>, 0.067 mg/mL diaphorase, 0.067 mM resazurin (in 50 mM Tris pH 8, 5 mM MgCl<sub>2</sub>, 0.01% Brij 35). |
| 2 | Reagent b | 3 μl | Dispense reagent b: 0.2 mM NAD<sup>+</sup>, 0.067 mg/mL diaphorase, 0.067 mM resazurin (in 50 mM Tris pH 8, 5 mM MgCl<sub>2</sub>, 0.01% Brij 35). |
| 3 | Compounds | 23 nL | Pin transfer of compounds |
| 4 | Time | 10 min | Incubate at room temperature |
| 5 | Reagent | 1 μl | Dispense 1 μl 0.16 mM malic acid |
| 6 | Read | 525 nm / 598 nm | ViewLux read twice at 5 minutes interval |

### qHTS screening and counter-screen

We screened six in-house libraries with a total number of 57,037 small molecule compounds in 1536-well format (Fig 2A). The libraries include L0PAC®1280 which is the Library of Pharmacologically Active Compounds, NCATS Pharmaceutical Collection which covers 2,816 compounds from late-stage clinical trials and FDA approved drugs, NCATS Pharmacologically Active Chemical Toolbox which contains 5,099 annotated compounds with known targets or biological pathways, MIPE (Mechanism Interrogation Plate) collection with 1,912 oncology focused compounds with well-annotated primary mechanisms of action, a kinase inhibitor focused library with 977 compounds, and a small molecule screening collection with diverse scaffolds designed from novel chemical space (44,953 compounds). The assay was robust and consistent across the screening plates, with an average assay Z’ factor 0.73 ± 0.09 (Fig 2B). The hit rate of activators was 0.1% (69 compounds), and inhibitors was 1.5% (868 compounds). The qHTS data was deposited in PubChem with AID 1347055, and you can access this assay with the following link: https://pubchem.ncbi.nlm.nih.gov/assay/assay.cgi?aid=1347055. We cherrypicked 374 compounds based on our selection criteria (active compounds and efficacy ≥ 50%, see methods) for confirmation testing. In the confirmation assay, 13 compounds were confirmed as active activators and 178 as active inhibitors. However, the majority of the confirmed hits displayed activity in the MDH counter-assay and were triaged.

**Figure 2.**
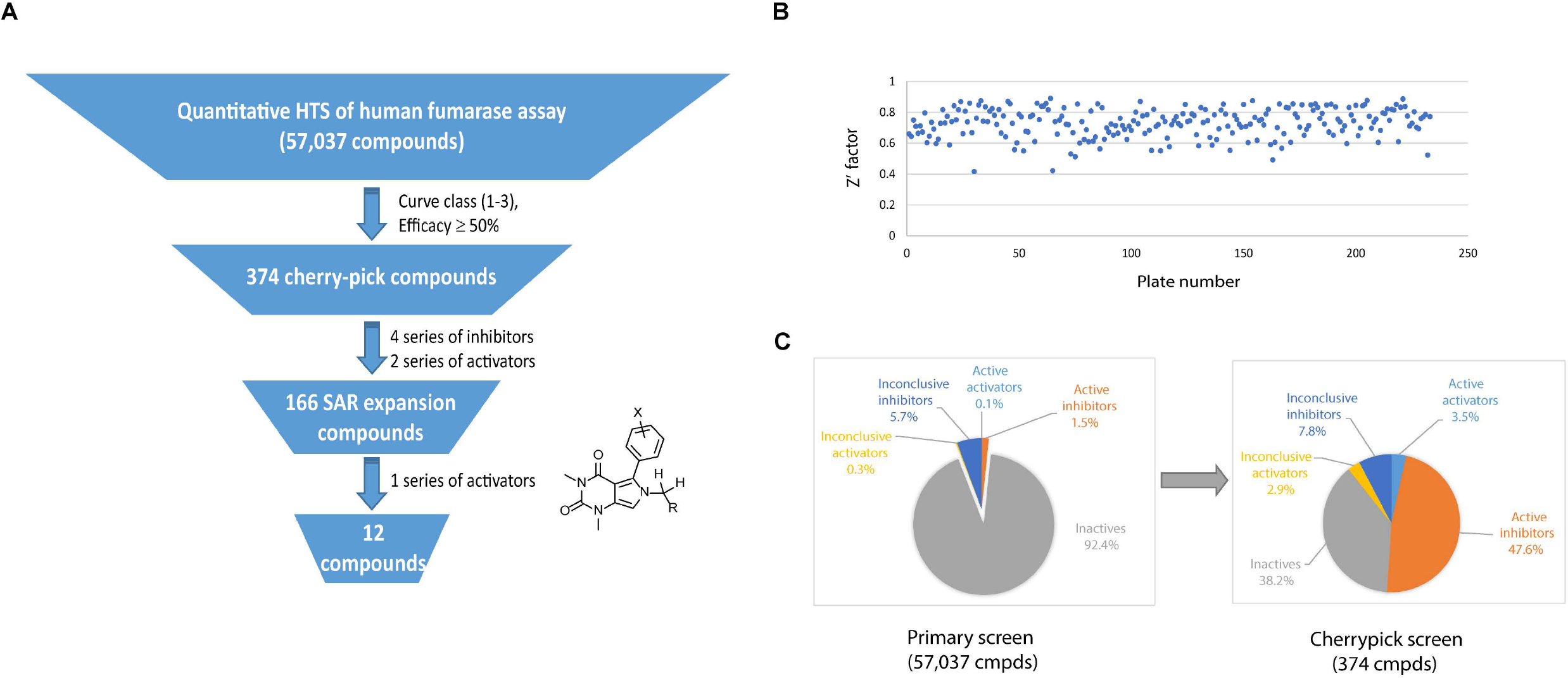
Statistics of quantitative high-throughput screen (qHTS) of human fumarase assay. **A**. Flow chat of qHTS of human fumarase assay. 57,037 compounds from six libraries were screened in the primary screen. 374 compounds, which belong to the curve class 1-3 and efficacy is more than 50%, were cherry-picked as hits. After confirmative assay, 166 compounds expanded from 4 series of inhibitors and 2 series of activators were picked for the further experiment. 12 compounds from one series of activators were confirmed and selected for the follow up experiments. **B**. Dot plot of Z factor of human fumarase assay from 233 primary screen plates. Z factor is ranged from 0.4 to 0.9 and average of Z factor is 0.73. **C**. Pie charts of hit rates of primary and confirmation screen of human fumarase assays. The compounds were classified as five categories: inactives (curve class 4), active activators (curve class 1.1, 1.2, 2.1, 2.2), active inhibitors (curve class −1.1, −1.2, −2.1, −2.2), inclusive activators (curve class 1.3, 1.4, 2.3, 2.4, 3) and inclusive inhibitors (curve class −1.3, −1.4, −2.3, −2.4, −3). The hit rate of primary screen is shown in left panel and the hit rate of confirmation screen is shown in right panel.

Only 4 series of inhibitors and 2 series of activators survived the counter-screening process. We then plated a total of 166 analogs structurally related to the hits from our in-house library for re-testing. One series of activators was confirmed in our FH assay (Fig 3A) and further evaluated in the follow up experiments. All other compounds dropped out during the reconfirmation assays.

**Figure 3.**
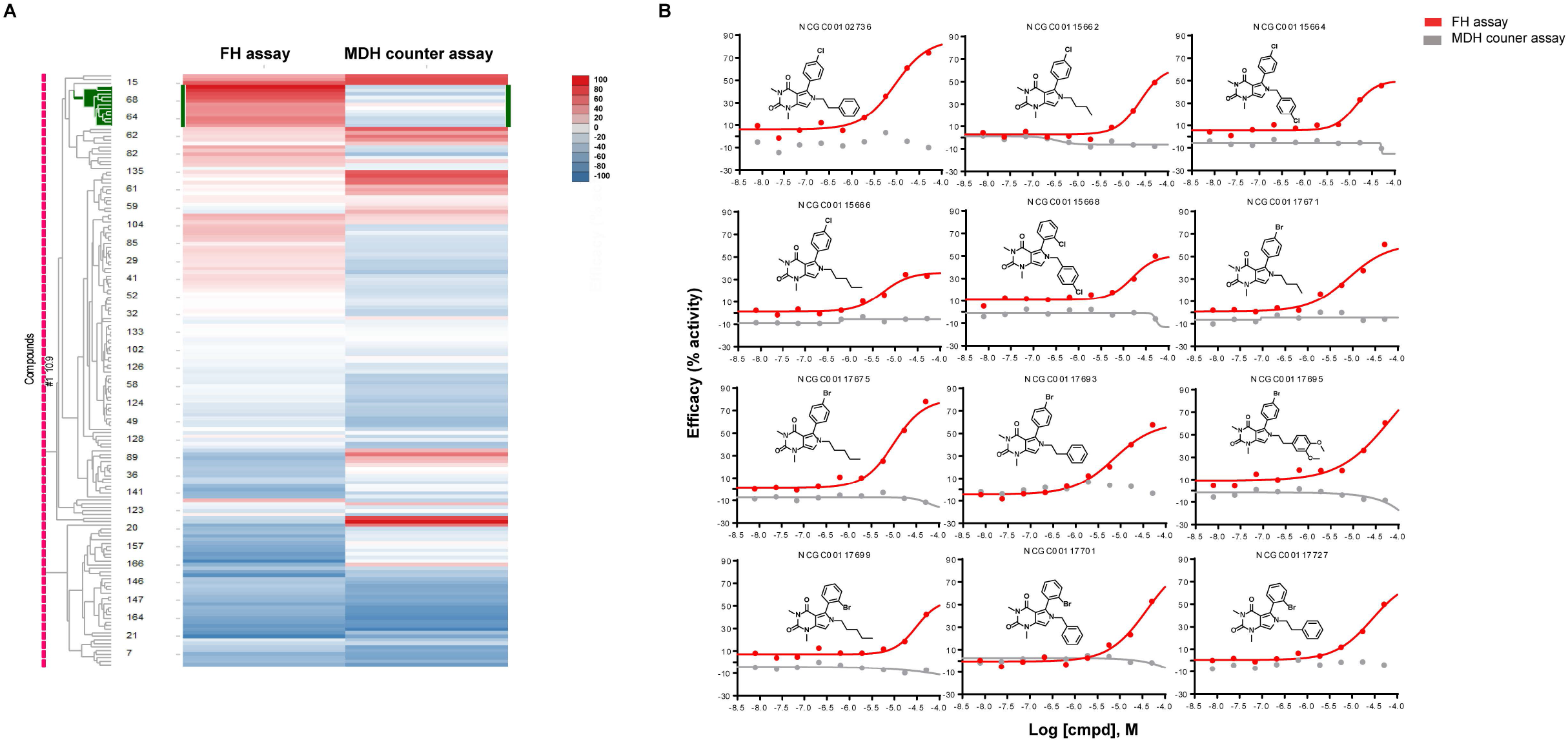
Heatmap of 166 compounds from six structural series. **A**. Hierarchical clustering analysis was done using Spotfire DecisionSite 8.2. In the heat map, each row represents a compound and each column represents an assay readout. The left column is fumarase assay and right column is malate dehydrogenase (MDH) assay. Each cluster of compounds is labelled by the most significantly enriched MeSH PA term in that cluster measured by a Fisher’s exact test. The color in heat map represent the activity of the compound, ranging from blue (inhibitors) to red (activators). **B**. Dose response curve of one series of activators (12 compounds). The X-axis is the l logarithmic scale of compound concentration and Y-axis is the efficacy (%activity) of the compounds. Red curves represent the data from FH assay and grey curves represent the data from MDH counter assay. NCGC number is the sample identification number of each compound in our library.

### Identification and validation of FH activators

The activator chemotype is composed of a series of phenyl-pyrrolo-pyrimidine-diones which are active against FH with AC_50_ ranging from 5.6 μM to 31.6 μM and efficacy ranging from 35% to 79% (Table 2). Compounds were not active in the MDH counter assay (Fig 3B). No signal was detected in the absence of substrates in both assays (Fig 4A), which indicated neither compound was the substrate of those two enzymes (unlike tartrate, vide infra). To ensure activity was not due to impurities in the library material, four compounds (highest potency and efficacy) were procured from vendors and purified via reverse phase HPLC. The newly acquired compounds showed similar activity (Supplemental Fig 2).

**Figure 4.**
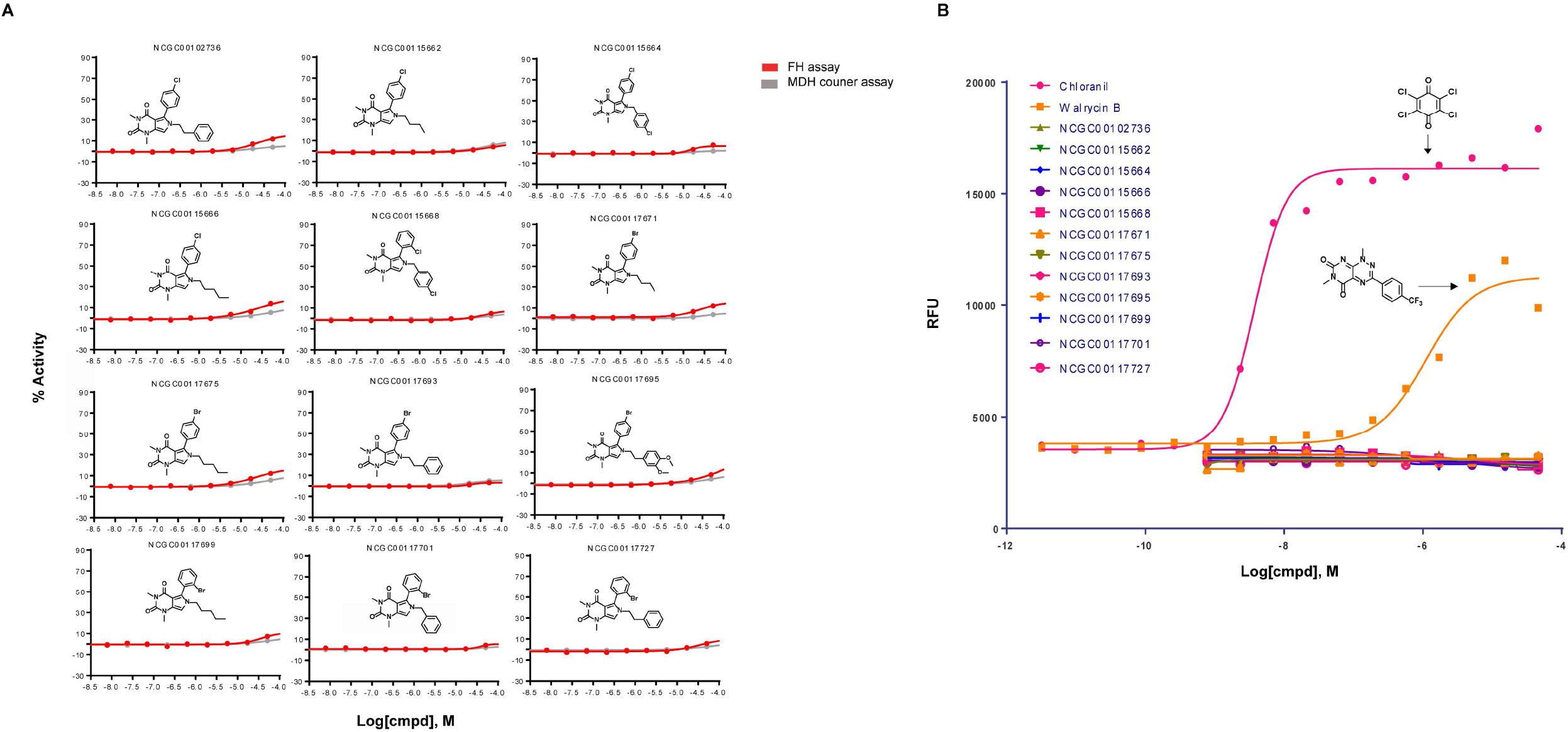
Evaluation of the series of activators on FH activity without fumarate and REDOX activity in Amplex assay. **A.** No FH activity of the series of activators were shown in the absence of fumarate (substrate of fumarase). The effect of the series of activators on FH or MDH activity was evaluated in FH assay or MDH assay without fumarate or malate respectively. The protocols were same except the substrates were omitted from the assays. Red curves represent the data from FH assay and grey curves represent the data from MDH counter assay. **B.** No reduction-oxidation reaction (REDOX) activity of the series of activators were detected in Amplex red assay. REDOX activity of activators was evaluated in Amplex red assay. Two known compounds with different REDOX activity were used as positive controls. Chloranil has a strong REDOX activity and walrycin B has a weak REDOX activity.

**Table 2.**
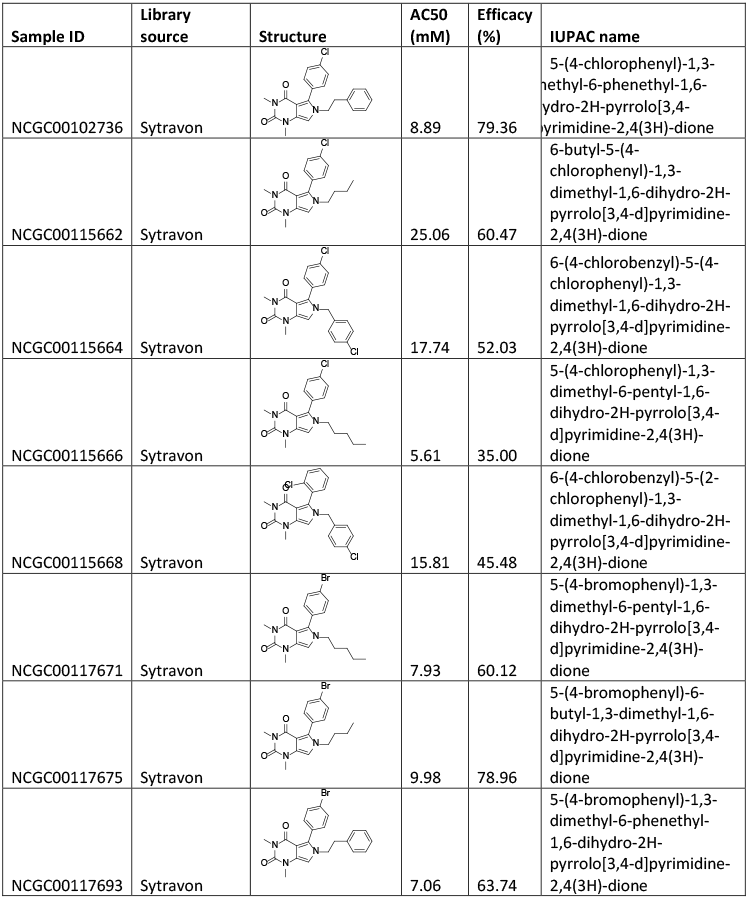
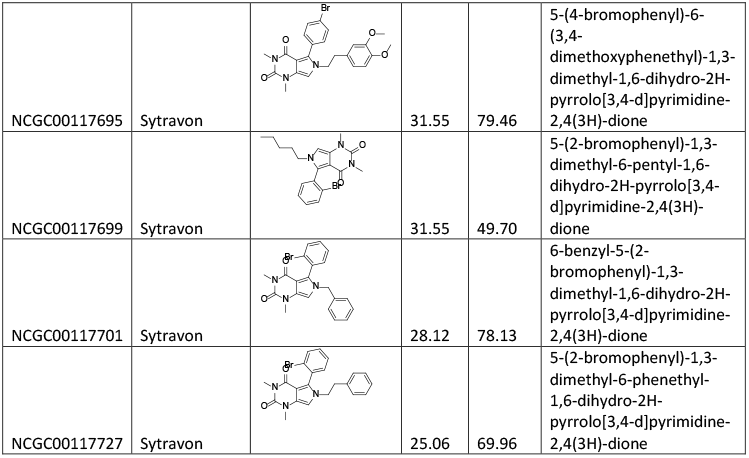
One series of activators of FH.

| Sample ID | Library source | Structure | AC50 (mM) | Efficacy (%) | IUPAC name |
| --- | --- | --- | --- | --- | --- |
| NCGC00102736 | Sytravon |  | 8.89 | 79.36 | 5-(4-chlorophenyl)-1,3-dimethyl-6-phenethyl-1,6-dihydro-2H-pyrrolo[3,4-d]pyrimidine-2,4(3H)-dione |
| NCGC00115662 | Sytravon |  | 25.06 | 60.47 | 6-butyl-5-(4-chlorophenyl)-1,3-dimethyl-1,6-dihydro-2H-pyrrolo[3,4-d]pyrimidine-2,4(3H)-dione |
| NCGC00115664 | Sytravon |  | 17.74 | 52.03 | 6-(4-chlorobenzyl)-5-(4-chlorophenyl)-1,3-dimethyl-1,6-dihydro-2H-pyrrolo[3,4-d]pyrimidine-2,4(3H)-dione |
| NCGC00115666 | Sytravon |  | 5.61 | 35.00 | 5-(4-chlorophenyl)-1,3-dimethyl-6-pentyl-1,6-dihydro-2H-pyrrolo[3,4-d]pyrimidine-2,4(3H)-dione |
| NCGC00115668 | Sytravon |  | 15.81 | 45.48 | 6-(4-chlorobenzyl)-5-(2-chlorophenyl)-1,3-dimethyl-1,6-dihydro-2H-pyrrolo[3,4-d]pyrimidine-2,4(3H)-dione |
| NCGC00117671 | Sytravon |  | 7.93 | 60.12 | 5-(4-bromophenyl)-1,3-dimethyl-6-pentyl-1,6-dihydro-2H-pyrrolo[3,4-d]pyrimidine-2,4(3H)-dione |
| NCGC00117675 | Sytravon |  | 9.98 | 78.96 | 5-(4-bromophenyl)-6-butyl-1,3-dimethyl-1,6-dihydro-2H-pyrrolo[3,4-d]pyrimidine-2,4(3H)-dione |
| NCGC00117693 | Sytravon |  | 7.06 | 63.74 | 5-(4-bromophenyl)-1,3-dimethyl-6-phenethyl-1,6-dihydro-2H-pyrrolo[3,4-d]pyrimidine-2,4(3H)-dione |
| NCGC00117695 | Sytravon |  | 31.55 | 79.46 | 5-(4-bromophenyl)-6-(3,4-dimethoxyphenethyl)-1,3-dimethyl-1,6-dihydro-2H-pyrrolo[3,4-d]pyrimidine-2,4(3H)-dione |
| NCGC00117699 | Sytravon |  | 31.55 | 49.70 | 5-(2-bromophenyl)-1,3-dimethyl-6-pentyl-1,6-dihydro-2H-pyrrolo[3,4-d]pyrimidine-2,4(3H)-dione |
| NCGC00117701 | Sytravon |  | 28.12 | 78.13 | 6-benzyl-5-(2-bromophenyl)-1,3-dimethyl-1,6-dihydro-2H-pyrrolo[3,4-d]pyrimidine-2,4(3H)-dione |
| NCGC00117727 | Sytravon |  | 25.06 | 69.96 | 5-(2-bromophenyl)-1,3-dimethyl-6-phenethyl-1,6-dihydro-2H-pyrrolo[3,4-d]pyrimidine-2,4(3H)-dione |

**Table 3.** *In vitro* ADME of top four compounds

| Sample ID | Rat liver microsome stability (t<sub>1/2</sub>, min) | PAMPA Permeability pH 7.4 (1x10<sup>-6</sup> cm/s) | PAMPA permeability pH 5 (1x10<sup>-6</sup> cm/s) | Solubility (ug/ml) | cLogP |
| --- | --- | --- | --- | --- | --- |
| NCGC00102736 | 1.11 | 1,224.00 | 762.26 | 2.15 | 3.81 |
| NCGC00117671 | 6.39 | Not conclusive | 945.87 | <1 | 4.08 |
| NCGC00117675 | 7.68 | 1,507 | 1,194 | <1 | 3.57 |
| NCGC00117693 | 1.23 | Not conclusive | 590 | 14.7 | 3.94 |

Artefactual liabilities of screening hits can be responsible for the apparent activation, such as the ability to undergo reduction-oxidation (redox) cycling. To examine whether compounds demonstrate redox activity, the amplex red assay was performed. In the assay, two compounds with weak (Walrycin B) and strong (Chloranil) redox activity were used as positive controls. None of the hit compounds displayed detectible redox activity (Fig 4B).

We performed microscale thermophoresis as a biophysical assay to directly demonstrate binding of activators to FH^21^. Two compounds with high potency, NCGC00102736 (AC_50_ = 8.89 μM) and NCGC00117675 (AC_50_ = 9.98 μM) were evaluated. The dissociation constant (K_d_) of NCGC00102736 was found to be 18.3 μM and NCGC00117675 was 23.3 μM (Fig 5).

**Figure 5.**
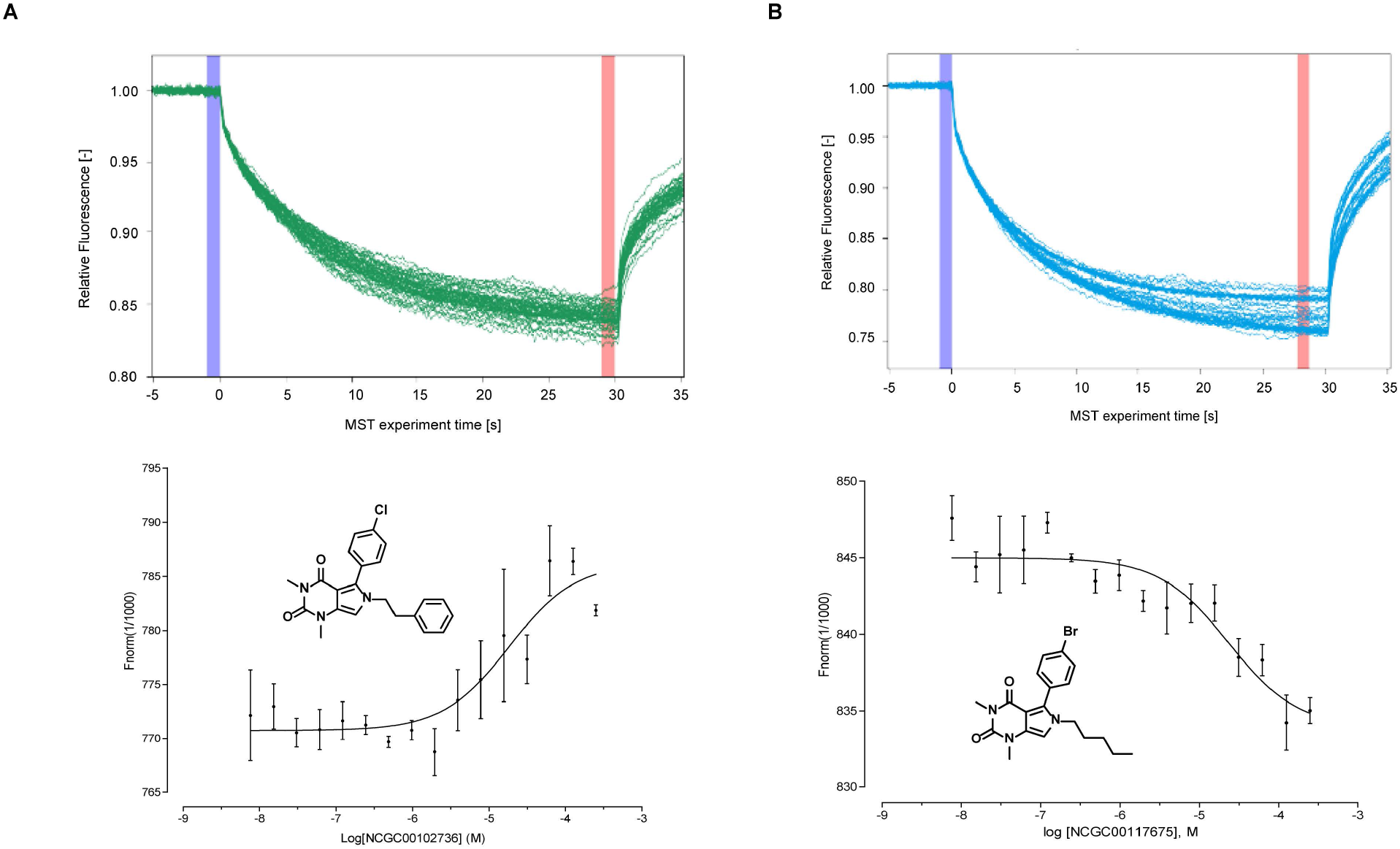
Target engagement of top two compounds with fumarate hydratase. The binding of two compounds with fumarase enzyme were evaluated by microscale themophoresis (MST) assay. *Top panel:* MST traces of titrations of NCGC00102736 (**A**) and NCGC00117675 (**B**) against 50 nM human Fumarate hydratase. The cold region (blue) and the hot region (red) were used to determine the Kd of the interaction of compounds and FH enzymes. *Low panel:* Dose-response curve for the binding interaction between compounds and FH enzymes. The binding curve yields a Kd of 18.3uM for NCGC00102736 (**A**) and 23.3uM for NCGC00117675 (**B**).

### *In vitro* ADME properties of FH activators

To evaluate the physicochemical and pharmacokinetic properties of the FH activator series, we selected the four most potent compounds and tested them in a series of *in vitro* ADME assays. Using automated high-throughput *in vitro* assays, we examined rat microsomal metabolic stability, permeability and solubility of those compounds. The half-life (t1/2) of compounds in rat hepatic microsomal stability assay ranged from 1.1 min to 7.7 min, suggesting that these compounds were metabolically unstable. The PAMPA was used as an *in vitro* model of passive, transcellular permeation. Two pH conditions (pH 5 and pH 7.5) were tested in this assay. All compounds scored as highly permeable (>100 x 10^6^ cm/s) at pH 5 and two compounds (NCGC00102736 and NCGC00117675) showed high permeability at pH 7.4. Aqueous solubility was assessed by a high-throughput kinetic solubility assay. The value of solubility ranged from <1 μg/mL to 14.7 μg/mL, indicating these compounds had low aqueous solubility.

### Evaluation of prior art compounds and artifacts

As two inhibitors of FH were reported previously, we evaluated the activity of those inhibitors in our human FH assay. We re-synthesized the compound 3 (fumarate hydratase-IN-2) which had been reported as active against human FH in a commercial *in vitro* assay (K_i_=4.5 μM)^11^. We found compound 3 to be inactive in our biochemical assay (Fig 6A). Based on the core structure of compound 3, we further cherrypicked 68 analogs of this chemotype and found none of them were active in our FH assay (data not shown). In another report, Kasbekar^12^ reported two inhibitors of the *Mycobacterium tuberculosis* fumarate hydratase (tbFH) from the same chemotype (compound 7 and 8 in their paper). We tested compound 8, which showed weak inhibitory activity at the highest concentration (IC_50_ = 42.2 μM and efficacy = 30.2%) in our human FH assay (Fig 6A). Given that enzymes from two species share identical residues in active sites and overall 59% similarity^22^, it is not surprising that compounds inhibitor targeting tbFH also showed a weak inhibitory activity against human FH.

**Figure 6.**
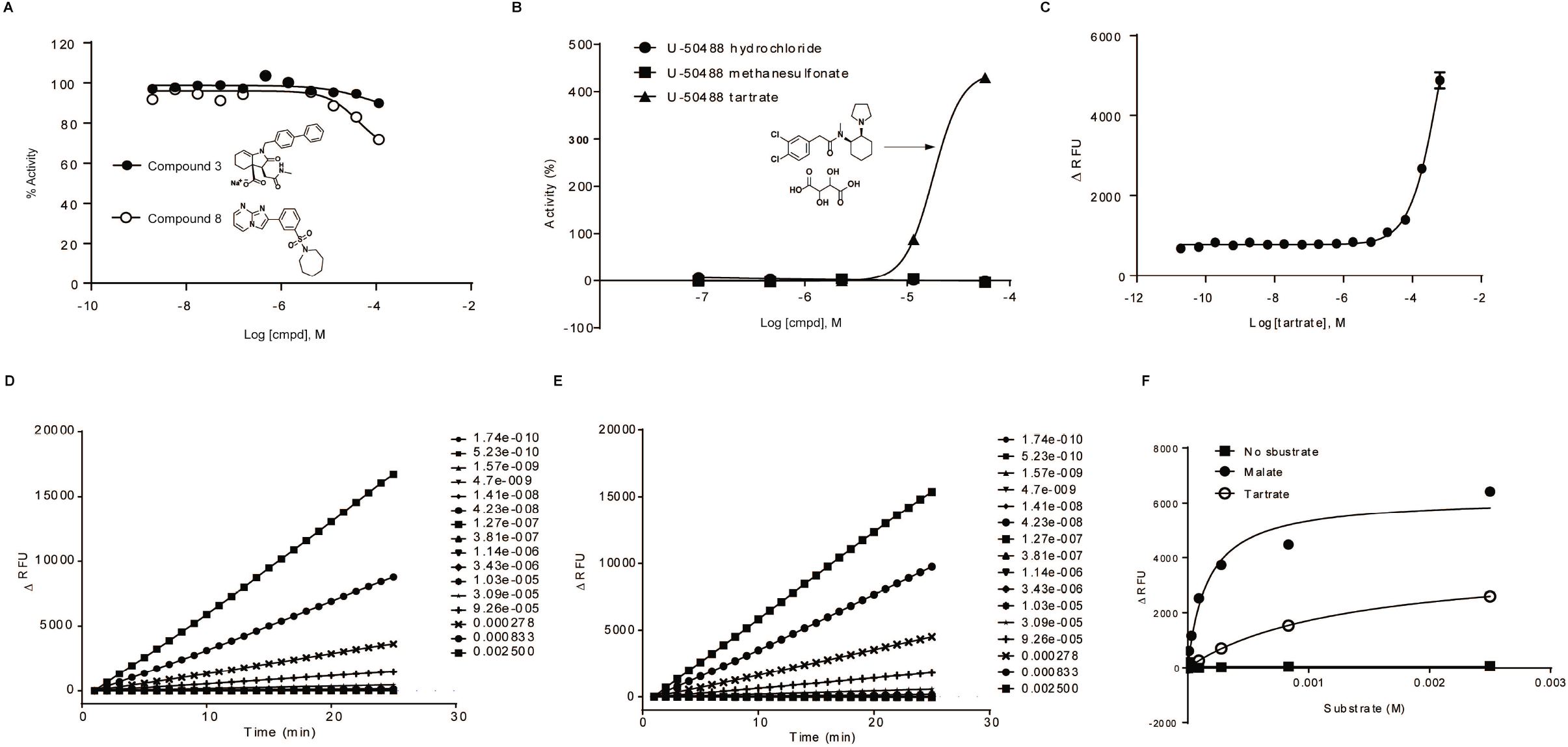
Evaluation of prior art compounds and artifacts. A. The reported compound 3 and compound 8 were resynthesized and tested in FH assay. **B**. Activity of different salt format of U-50488 compound in FH assay. **C**. Activity of tartrate salt in FH assay. **D**. Time course of FH assay using tartrate as a substrate for the reaction without adding fumarate. **E**. Time course of MDH assay using tartrate as a substrate for the reaction without adding malate. **F**. Determine Km value of tartrate for MDH enzyme.

During our HTS campaign, we identified an interesting false-positive compound, U-50488, which is a highly selective κ-opioid agonist. There are three salt forms of U-50488 in our collection. We found that the tartrate salt of U-50488 showed very high activator activity. However, the hydrochloride and methanesulfonate salts of U-50488, also in our library, showed no activator activity (Fig 6B). We speculated that this activator activity might derive from the tartrate salt component, instead of compound U-50488 itself. Potassium sodium tartrate was then tested and was confirmed to produce increased activity in our FH assay (Fig 6C). As tartrate was reported as a substrate of FH and MDH in the 1960s^23, 24^, we hypothesized that tartrate might be utilized as the substrate of FH or MDH in our assays, giving the appearance of activation. To test this hypothesis, we serially diluted tartrate and added to FH or MDH assays without their normal substrate (fumarate or malate) in the reaction. We found that tartrate can drive both enzymes in the absence of their eponymous substrates with a similar reaction rate (Fig 6D and 6E). The K_m_ value of tartrate was determined by MDH assay with K_m_ = 1296 ± 33 μM (Fig 6F), much lower than malate, the normal substrate of MDH (K_m_ = 225 ± 30 μM).

## Discussion

In our efforts to identify the small molecule probes for FH, we developed a quantitative high-throughput screening (qHTS) FH assay and screened a collection of small molecule libraries. After several rounds of screening and counter screening, only one series of activators met all activity criteria in our study. Those compounds were not a substrate of FH, and they demonstrated direct binding to FH protein. This activator series enhanced the FH activity in the presence of its substrate. Our results suggest that these compounds are allosteric modulators of fumarate hydratase, however, further work is needed to optimize activity and confirm their mechanism of activation.

The series identified as FH activators is a group of phenyl-pyrrolo-pyrimidine-diones (Fig 2A and Fig 3B). The hit series consists of 12 active compounds and 9 inactive compounds. A structure-activity relationship (SAR) analysis with this limited set of compounds suggested that the nature of the substitution on the pyrrole nitrogen (R) was important for FH activation (Supplemental Table 1). Active compounds appeared to favor the attachment of a methylene unit to the pyrrole nitrogen followed by further substitution (see Figure 2A). Attachment of additional substituents to this carbon, for example when R was branched such as a phenyl ring or a secondary or tertiary alkyl group, led to erosion in activity. Linear alkyl, phenethyl, and benzyl substituents on the pyrrole N were well tolerated. These data suggest that the pyrrole nitrogen might be important for the compound’s interaction with the protein target and that an uncrowded environment around it might be essential for its binding to the target. With regard to the 5-phenyl ring the presence of a halide substitution (X = Cl or Br) appeared to be necessary for activity (Supplemental Table 1) as a compound (NCGC00141286) with a long alkyl R but no phenyl ring substitution was inactive. With only one exception, all active compounds in the series have a halogen in the para or ortho position. This information and medicinal chemistry efforts could further improve the chemotype’s potency against fumarase.

Pan-assay interference compounds (PAINS) are the compounds shown as false positives in high-throughput screens^25^. To exclude fluorescent compounds, we used change of fluorescence (ΔRFU) as the readout. If compounds are fluorescent, the signal should not change over the time (15 minutes in our assay) and the value of ΔRFU should be close to zero. To filter out the compounds interacting with coupled enzymes in the assay (MDH and diaphorase), a MDH assay was used as a counter-assay to triage these false-positive compounds. However, the MDH counter-assay may triaged compounds that inhibit or activate both FH and MDH. As a common substrate for both FH and MDH, malate should be able to bind to the active sites of both enzymes, which suggests a structural similarity in active sites between two enzymes. The inhibitors which competitively bind to same active sites will be filtered out in MDH counter assay. As such, non-selective competitive inhibitors would be prone to exclusion by assay format. No inhibitors were identified in current study after screening a collection of 57,037 compounds. In a similar assay format, Kasbekar^12^ also didn’t identify any competitive inhibitors for tbFH after screening a collection of 479,984 compound. Instead, two inhibitors that bind to an allosteric regulatory site were found.

In summary, we developed a quantitative high-throughput screening (qHTS) FH assay and screened a collection of 57,037 small molecules. A series of phenyl-pyrrolo-pyrimidine-diones were identified as activators of human fumarate hydratase, which can serve as a starting point for further optimization and development as probes for the enzyme.

## Acknowledgement

This work was supported by the intramural research program of the National Center for Advancing Translational Sciences (NCATS). We would thank Monica Kasbekar and Craig J. Thomas for kindly providing the detailed protocol of their fumarase assay; Paul Shinn, Danielle van Leer and Zina Itkin for the assistance with compound management; Sam Michael, Carleen Klumpp-Thomas and Jameson Travers for the assistance of automation; Michael Ronzetti and Bolormaa Baljinnyam for the technical support of microscale thermophoresis assay; Mark Henderson for the technical support of amplex red assay.

---

**Supplemental figure 1.** Thermostability of human FH incubated at room temperature or on ice. FH solutions containing human fumarate hydratase were incubated at RT (**A**) or on ice (**B**) at different time point (0hr, 2hr, 4hr, 6hr, 8hr and 24hr). The reaction was triggered by adding fumarate and fluorescence was continuously measured every minute for 20 minutes.

**Supplemental figure 2**. Validation of four compounds (highest potency and efficacy) which were procured from vendors and purified. Activity of these compounds were confirmed in FH assay.

## Reference

1. Weaver, T. M.; Levitt, D. G.; Donnelly, M. I.; et al. The multisubunit active site of fumarase C from Escherichia coli. Nat Struct Biol 1995, 2, 654–62.

2. Picaud, S.; Kavanagh, K. L.; Yue, W. W.; et al. Structural basis of fumarate hydratase deficiency. J Inherit Metab Dis 2011, 34, 671–6.

3. Ewbank, C.; Kerrigan, J. F.; Aleck, K. Fumarate Hydratase Deficiency. In GeneReviews((R)); Adam, M. P.; Ardinger, H. H.; Pagon, R. A.; et al., Eds.; Seattle (WA), 1993.

4. Bourgeron, T.; Chretien, D.; Poggi-Bach, J.; et al. Mutation of the fumarase gene in two siblings with progressive encephalopathy and fumarase deficiency. J Clin Invest 1994, 93, 2514–8.

5. Tomlinson, I. P.; Alam, N. A.; Rowan, A. J.; et al. Germline mutations in FH predispose to dominantly inherited uterine fibroids, skin leiomyomata and papillary renal cell cancer. Nat Genet 2002, 30, 406–10.

6. Yang, M.; Pollard, P. J. Succinate: a new epigenetic hacker. Cancer Cell 2013, 23, 709–11.

7. Yang, M.; Soga, T.; Pollard, P. J. Oncometabolites: linking altered metabolism with cancer. J Clin Invest 2013, 123, 3652–8.

8. Tian, Z.; Liu, Y.; Usa, K.; et al. Novel role of fumarate metabolism in dahl-salt sensitive hypertension. Hypertension 2009, 54, 255–60.

9. You, Y. H.; Quach, T.; Saito, R.; et al. Metabolomics Reveals a Key Role for Fumarate in Mediating the Effects of NADPH Oxidase 4 in Diabetic Kidney Disease. J Am Soc Nephrol 2016, 27, 466–81.

10. Adam, J.; Ramracheya, R.; Chibalina, M. V.; et al. Fumarate Hydratase Deletion in Pancreatic beta Cells Leads to Progressive Diabetes. Cell Rep 2017, 20, 3135–3148.

11. Takeuchi, T.; Schumacker, P. T.; Kozmin, S. A. Identification of fumarate hydratase inhibitors with nutrient-dependent cytotoxicity. J Am Chem Soc 2015, 137, 564–7.

12. Kasbekar, M.; Fischer, G.; Mott, B. T.; et al. Selective small molecule inhibitor of the Mycobacterium tuberculosis fumarate hydratase reveals an allosteric regulatory site. Proc Natl Acad Sci U S A 2016, 113, 7503–8.

13. Huang, R.; Southall, N.; Wang, Y.; et al. The NCGC pharmaceutical collection: a comprehensive resource of clinically approved drugs enabling repurposing and chemical genomics. Sci Transl Med 2011, 3, 80ps16.

14. Urban, D. J.; Martinez, N. J.; Davis, M. I.; et al. Assessing inhibitors of mutant isocitrate dehydrogenase using a suite of pre-clinical discovery assays. Sci Rep 2017, 7, 12758.

15. Inglese, J.; Auld, D. S.; Jadhav, A.; et al. Quantitative high-throughput screening: a titration-based approach that efficiently identifies biological activities in large chemical libraries. Proc Natl Acad Sci U S A 2006, 103, 11473–8.

16. Wang, Y.; Huang, R. Correction of Microplate Data from High Throughput Screening. In High-Throughput Screening Assays in Toxicology; 1 ed.; Zhu, H.; Xia, M., Eds.; Methods in Molecular Biology; Humana Press: 2016; Chapter 13.

17. Hill, A. V. The possible effects of the aggregation of the molecules of haemoglobin on its dissociation curves. J. Physiol. (London) 1910, 40, 4–7.

18. Huang, R.; Xia, M.; Cho, M. H.; et al. Chemical genomics profiling of environmental chemical modulation of human nuclear receptors. Environ Health Perspect 2011, 119, 1142–8.

19. Davis, M. I.; Shen, M.; Simeonov, A.; et al. Diaphorase Coupling Protocols for Red-Shifting Dehydrogenase Assays. Assay Drug Dev Technol 2016, 14, 207–12.

20. Hall, M. D.; Simeonov, A.; Davis, M. I. Avoiding Fluorescence Assay Interference-The Case for Diaphorase. Assay Drug Dev Technol 2016, 14, 175–9.

21. Bartoschik, T.; Galinec, S.; Kleusch, C.; et al. Near-native, site-specific and purification-free protein labeling for quantitative protein interaction analysis by MicroScale Thermophoresis. Sci Rep 2018, 8, 4977.

22. Estevez, M.; Skarda, J.; Spencer, J.; et al. X-ray crystallographic and kinetic correlation of a clinically observed human fumarase mutation. Protein Sci 2002, 11, 1552–7.

23. Tsukatani, T.; Matsumoto, K. Enzymatic Quantification of L-Tartrate in Wines and Grapes by Using the Secondary Activity of D-Malate Dehydrogenase. Biosci Biotechnol Biochem 1999, 63, 1730–5.

24. Nakamura, S.; Ogata, H. Specificity of fumarate hydratase. I. Formation of oxalacetate from unnatural (--)-tartrate by fumarate hydratase. J Biol Chem 1968, 243, 528–32.

25. Baell, J. B.; Nissink, J. W. M. Seven Year Itch: Pan-Assay Interference Compounds (PAINS) in 2017-Utility and Limitations. ACS Chem Biol 2018, 13, 36–44.

