## Supplemental info for "Identification of activators of human fumarate hydratase by quantitative high-throughput screening"

### Supplemental figure 1

A B

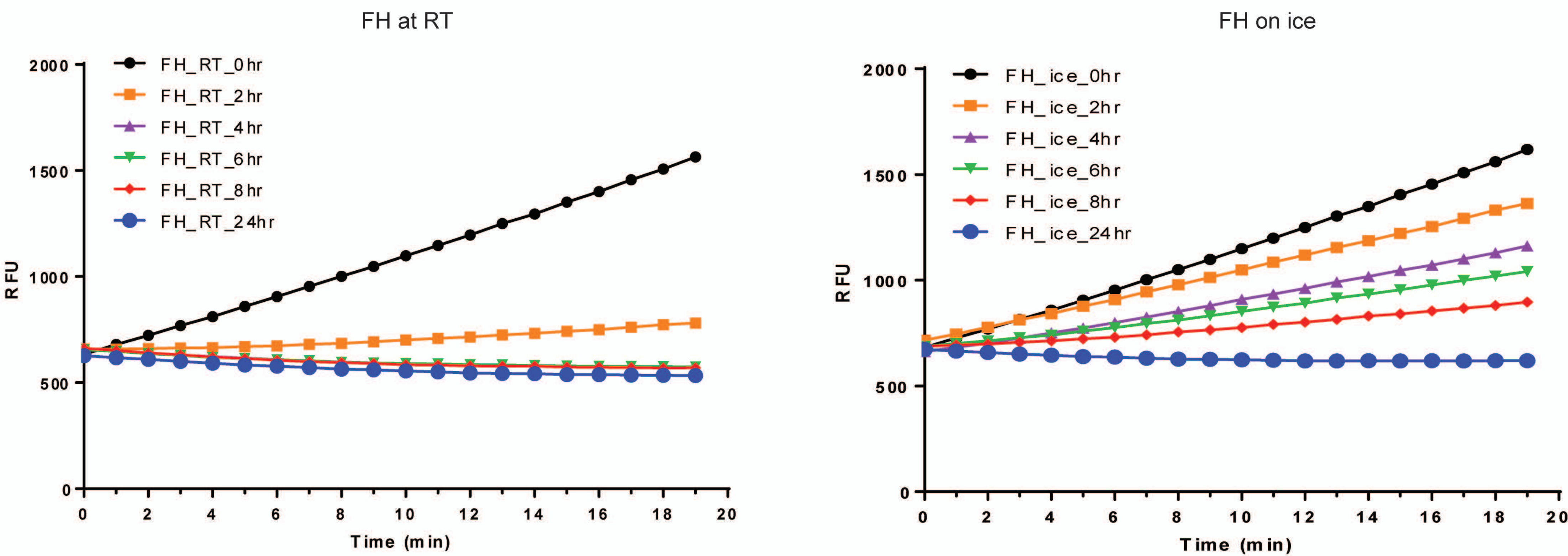

### Supplemental figure 2

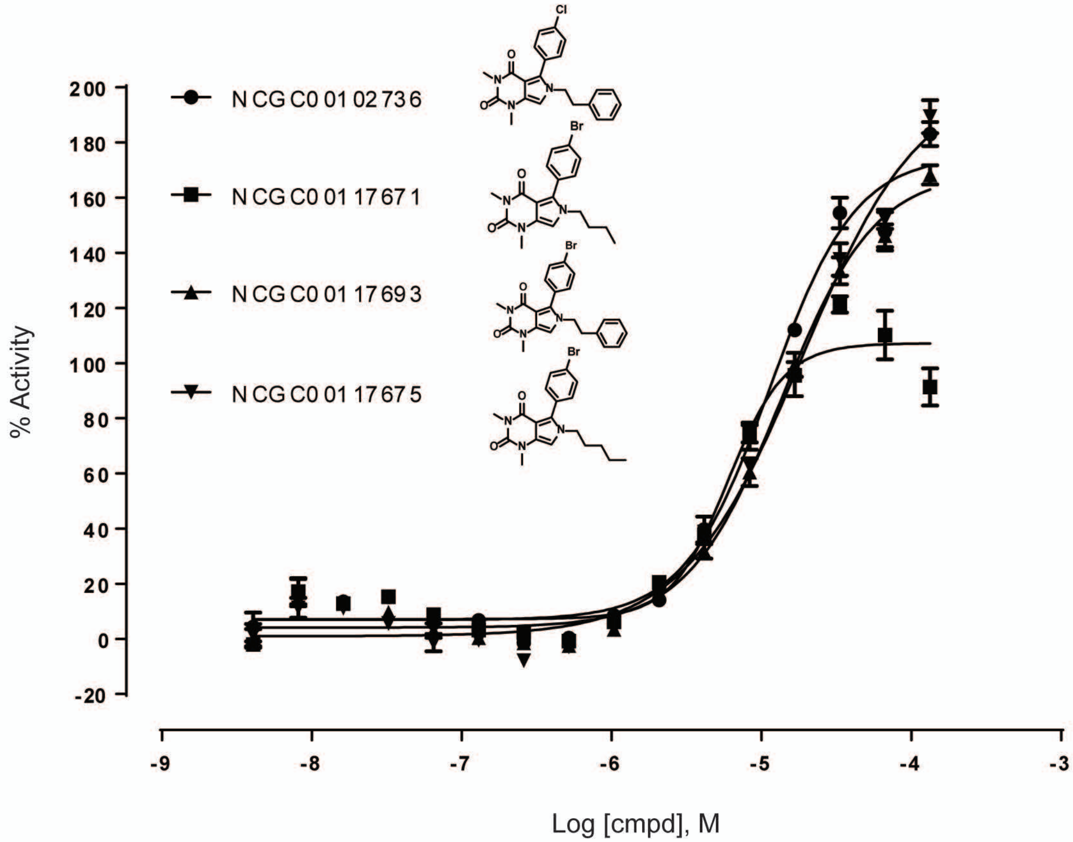

**Supplemental Table 1. Expansion of the series of activators of FH**

| Sample ID | Library source | Structure | AC50 (uM) | Efficacy (%) | Smiles |
|---|---|---|---|---|---|
| NCGC00102736 | Sytravon | 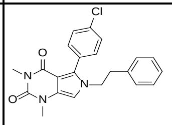 | 8.893 | 79 | CN1C(=O)N(C)C2=C[N](CCC3=CC=CC=C3)C(=C2C1=O)C4=CC=C(Cl)C=C4 |
| NCGC00115662 | Sytravon | 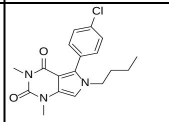 | 25.064 | 60 | CCCC[N]1C=C2N(C)C(=O)N(C)C(=O)C2=C1C3=CC=C(Cl)C=C3 |
| NCGC00115664 | Sytravon | 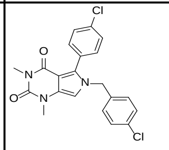 | 17.744 | 52 | CN1C(=O)N(C)C2=C[N](CC3=CC=C(Cl)C=C3)C(=C2C1=O)C4=CC=C(Cl)C=C4 |
| NCGC00115666 | Sytravon | 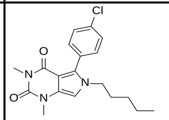 | 5.611 | 35 | CCCCC[N]1C=C2N(C)C(=O)N(C)C(=O)C2=C1C3=CC=C(Cl)C=C3 |
| NCGC00115668 | Sytravon | 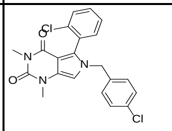 | 15.814 | 45 | CN1C(=O)N(C)C2=C[N](CC3=CC=C(Cl)C=C3)C(=C2C1=O)C4=C(Cl)C=CC=C4 |
| NCGC00115672 | Sytravon | 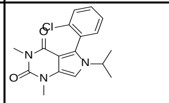 | null | 0 | CC(C)[N]1C=C2N(C)C(=O)N(C)C(=O)C2=C1C3=C(Cl)C=CC=C3 |
| NCGC00117671 | Sytravon | 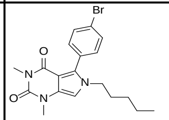 | 7.926 | 60 | CCCCC[N]1C=C2N(C)C(=O)N(C)C(=O)C2=C1C3=CC=C(Br)C=C3 |
| NCGC00117673 | Sytravon | 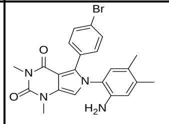 | 111.956 | -18 | CN1C(=O)N(C)C2=C[N](C3=C(N)C=C(C)C(=C3)C)C(=C2C1=O)C4=CC=C(Br)C=C4 |
| NCGC00117675 | Sytravon | 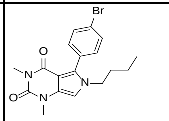 | 9.978 | 79 | CCCC[N]1C=C2N(C)C(=O)N(C)C(=O)C2=C1C3=CC=C(Br)C=C3 |
| NCGC00117693 | Sytravon | 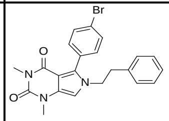 | 7.064 | 64 | CN1C(=O)N(C)C2=C[N](CCC3=CC=CC=C3)C(=C2C1=O)C4=CC=C(Br)C=C4 |
| NCGC00117695 | Sytravon | 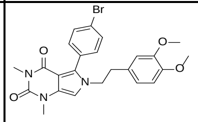 | 31.553 | 79 | COC1=CC=C(CC[N]2C=C3N(C)C(=O)N(C)C(=O)C3=C2C4=CC=C(Br)C=C4)C=C1OC |

| | | | | | |
|---|---|---|---|---|---|
| NCGC00117697 | Sytravon | 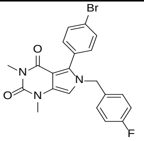 | 0.018 | 14 | CN1C(=O)N(C)C2=C[N](CC3=CC=C(F)C=C3)C(=C2C1=O)C4=CC=C(Br)C=C4 |
| NCGC00117699 | Sytravon | 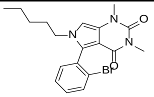 | 31.553 | 50 | CCCCC[N]1C=C2N(C)C(=O)N(C)C(=O)C2=C1C3=C(Br)C=CC=C3 |
| NCGC00117701 | Sytravon | 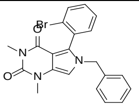 | 28.122 | 78 | CN1C(=O)N(C)C2=C[N](CC3=CC=CC=C3)C(=C2C1=O)C4=C(Br)C=CC=C4 |
| NCGC00117703 | Sytravon | 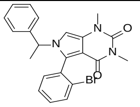 | 39.723 | 30 | CC([N]1C=C2N(C)C(=O)N(C)C(=O)C2=C1C3=CC=CC=C3Br)C4=CC=CC=C4 |
| NCGC00117707 | Sytravon | 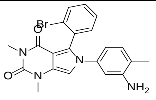 | 99.781 | -16 | CN1C(=O)N(C)C2=C[N](C3=CC=C(C)C(=C3)N)C(=C2C1=O)C4=CC=CC=C4Br |
| NCGC00117725 | Sytravon | 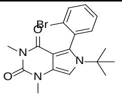 | 50.009 | 9 | CN1C(=O)N(C)C2=C[N](C(=C2C1=O)C3=CC=CC=C3Br)C(C)(C)C |
| NCGC00117727 | Sytravon | 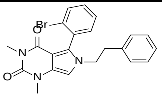 | 25.064 | 70 | CN1C(=O)N(C)C2=C[N](CCC3=CC=CC=C3)C(=C2C1=O)C4=CC=CC=C4Br |
| NCGC00117729 | Sytravon | 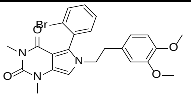 | 50.009 | 24 | COC1=C(OC)C=C(CC[N]2C=C3N(C)C(=O)N(C)C(=O)C3=C2C4=CC=CC=C4Br)C=C1 |
| NCGC00119865 | Sytravon | 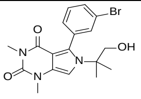 | null | 0 | CN1C(=O)N(C)C2=C[N](C(=C2C1=O)C3=CC(=CC=C3)Br)C(C)(C)CO |
| NCGC00141286 | Sytravon | 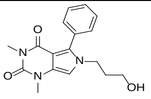 | 0.007 | -19 | CN1C2=CN(CCCO)C(=C2C(=O)N(C)C1=O)C3=CC=CC=C3 |
